# The critical immunosuppressive effect of MDSC-derived exosomes in the tumor microenvironment

**DOI:** 10.1101/2020.03.05.979195

**Authors:** Mohammad H. Rashid, Thaiz F. Borin, Roxan Ara, Raziye Piranlioglu, Bhagelu R. Achyut, Hasan Korkaya, Yutao Liu, Ali S. Arbab

**Affiliations:** Laboratory of Tumor Angiogenesis, Georgia Cancer Center, Department of Biochemistry and Molecular Biology, Augusta University, Augusta, Georgia, USA; Nanomedicine Research Center, Department of Neurosurgery, Cedars-Sinai Medical Center, Los Angeles, California, USA; Cancer Animal Models Shared Resource, Winship Cancer Institute, Emory University, Atlanta, Georgia, USA; Molecular Oncology and Biomarkers Program, Georgia Cancer Center, Department of Biochemistry and Molecular Biology, Augusta University, Augusta, Georgia, USA; Department of Cellular Biology and Anatomy, Medical College of Georgia, Augusta University, Augusta, Georgia, USA

**Author notes:** Corresponding author: Ali S. Arbab, MD, PhD, Georgia Cancer Center, Augusta University, 1410 Laney Walker Blvd, Room: CN 3315, Augusta, GA 30912.

**Keywords:** Exosomes, Myeloid-derived suppressor cells (MDSCs), Tumor microenvironment (TME), CD8 T-cell, activation-induced cell death (AICD)

## Abstract

Myeloid-derived suppressor cells (MDSCs) are an indispensable component of the tumor microenvironment (TME), and our perception regarding the role of MDSCs in tumor promotion is attaining extra layer of intricacy in every study. In conjunction with MDSC’s immunosuppressive and anti-tumor immunity, they candidly facilitate tumor growth, differentiation, and metastasis in several ways that yet to be explored. Alike any other cell types, MDSCs also release a tremendous amount of exosomes or nanovesicles of endosomal origin and partake in intercellular communications by dispatching biological macromolecules. There has not been any experimental study done to characterize the role of MDSCs derived exosomes (MDSC exo) in the modulation of TME. In this study, we isolated MDSC exo and demonstrated that they carry a significant amount of proteins that play an indispensable role in tumor growth, invasion, angiogenesis, and immunomodulation. We observed higher yield and more substantial immunosuppressive potential of exosomes isolated from MDSCs in the primary tumor area than those are in the spleen or bone marrow. Our *in vitro* data suggest that MDSC exo are capable of hyper activating or exhausting CD8 T-cells and induce reactive oxygen species production that elicits activation-induced cell death. We confirmed the depletion of CD8 T-cells *in vivo* by treating the mice with MDSC exo. We also observed a reduction in pro-inflammatory M1-macrophages in the spleen of those animals. Our results indicate that immunosuppressive and tumor-promoting functions of MDSC are also implemented by MDSC-derived exosomes which would open up a new avenue of MDSC research and MDSC-targeted therapy.

## Introduction

Apart from cancer cells, the tumor microenvironment (TME) accommodates heterogeneous host cells of the immune system, the tumor vasculature and lymphatics, fibroblasts, pericytes, and sometimes adipocytes [1]. Myeloid-derived suppressor cells (MDSCs) are crucial components of TME that play a pivotal role in tumor growth, neovascularization and metastasis [2, 3]. MDSCs are a group of vastly heterogeneous immunosuppressive cells derived from immature myeloid progenitors that have been linked to poor patient prognosis [4]. Typically immature myeloid cells traverse to the peripheral organs after originating from bone marrow and rapidly mature into macrophages, dendritic cells, or granulocytes (Neutrophils, Eosinophils, and Basophils) [5]. That said, in tumor condition, multifarious factors that are present in TME prevent the differentiation of these immature myeloid cells and instigate their actuation into an immunosuppressive phenotype [6]. MDSCs are usually divided into two subpopulations: gMDSCs (granulocytic, CD11b^+^Ly6G^+^Ly6C^low^), which are identical to neutrophils, and mMDSCs (monocytic, CD11b^+^Ly6G^−^Ly6C^hi^), which are consistent to monocytes in respect of morphology and phenotype [7, 8].

There is growing evidence that MDSCs harness various immune and nonimmune mechanisms to promote tumor development. MDSC inhibit adaptive antitumor immunity by inhibiting T cell activation and function (T-cell receptor downregulation, T-cell cycle inhibition and immune checkpoint blockade) [8], and by driving and recruiting T regulatory cells. Immunosuppression by MDSCs is also mediated by the generation of reactive oxygen species (ROS) [9] and cytokine release (IL-10, TGF-β) [10, 11], in conjunction with arginine depletion [12]. They inhibit innate immunity by polarizing macrophages toward a type 2 tumor-promoting phenotype (M2-macrophage) [13] and by inhibiting NK cell-mediated cytotoxicity [14]. Over and above MDSCs are efficient recruiters of other immunosuppressive cells.

Although the role of MDSCs in tumor growth and metastasis is well known, there is a significant knowledge gap for understanding the similar role of MDSC-derived exosomes (MDSC exo). In the past decade, there is a huge surge of exosome research and publications that are mostly focused on exosomes derived from tumor cells and immune cells. Exosomes are 30–150 nm lipid bi-layered extracellular bioactive vesicles of endosomal origin that are secreted by all the cells and present in various body fluids. Exosomes have been proposed to act as intercellular communicators as they can transfer their cargo (proteins, lipids, and nucleic acids) to nearby or distant recipient cells. Previously, we observed MDSC exo carry a significant amount of pro-tumorigenic factors, and a tremendous amount of MDSC exo injected intravenously was distributed in primary breast tumor and metastatic site [15]. These findings warrant us to further explore the implication of MDSC exo in immunosuppression and tumor progression mechanisms.

In this study, we characterized the size, yield, and contents of exosomes collected from different MDSC populations and immature myeloid progenitor cells. We now report that alike parental MDSCs, exosomes from the MDSCs also play a crucial role in inciting immunosuppressive milieu by way of limiting the functions of cytotoxic T cells and pro-inflammatory M1 macrophages in the TME.

## Materials and methods

### Ethics statement

All the experiments were performed according to the National Institutes of Health (NIH) guidelines and regulations. The Institutional Animal Care and Use Committee (IACUC) of Augusta University (protocol #2014–0625) approved all the experimental procedures. All animals were kept under regular barrier conditions at room temperature with exposure to light for 12 hours and dark for 12 hours. Food and water were offered ad libitum. All efforts were made to ameliorate the suffering of animals. CO2 with a secondary method was used to euthanize animals for tissue collection.

### Nanoparticle tracking analysis

Nanoparticle tracking analysis (NTA) was performed using ZetaView, a second-generation particle size instrument from Particle Metrix for individual exosome particle tracking as described preciously [15]. This is a high-performance integrated instrument equipped with a cell channel, which is integrated into a ‘slide-in’ cassette and a 405-nm laser. Samples were diluted in 1X PBS between 1:100 and 1:2000 and injected in the sample chamber with sterile syringes (BD Discardit II, New Jersey, USA). All measurements were performed at 23°C and pH 7.4. As measurement mode, we used 11 positions with 2 cycles, and for analysis parameter, we used maximum pixel 200 and minimum 5. ZetaView 8.02.31 software and Camera 0.703 μm/px were used for capturing and analyzing the data.

### Flow cytometry

For the *in vivo* flow cytometric analysis, the collected fresh tissue was disseminated into single cells, filtered through a 70 µm cell strainer, and spun at 1,200 rpm for 15 minutes. For the *in vitro* flow cytometric analysis, cells were washed twice with sterile PBS. The pellet was re-suspended in 1% BSA/PBS and incubated with LEAF blocker in 100 µL volume for 15 minutes on ice to reduce non-specific staining. The single cells were then labeled to detect the immune cell populations using fluorescence conjugated antibodies such as CD3, CD4, CD8, CD206, F4/80, CD279, CD25, CD184, CD194, CD69, CD62L, CD11b, CD80, CD86, Gr1, Ly6C, Ly6G and CD45. All antibodies were mouse-specific (Biolegend, San Diego, CA, USA) and the samples were acquired using the Accuri C6 flow cytometer (BD Biosciences).

### Tumor model

Both 4T1 and AT3 cells expressing the luciferase gene were orthotopically implanted in syngeneic BALB/c and C57BL/J6 mice, respectively (Jackson Laboratory, Main USA). All the mice were between 5-6 weeks of age and weighing 18-20g. Animals were anesthetized using a mixture of Xylazine (20mg/Kg) and Ketamine (100mg/Kg) administered intraperitoneally. Hair was removed for the right half of the abdomen by using hair removal ointment, and then the abdomen was cleaned by Povidone-iodine and alcohol. A small incision was made in the middle of the abdomen, and the skin was separated from the peritoneum using blunt forceps. Separated skin was pulled to the right side to expose the mammary fat pad and either 50,000 4T1 cells or 100,000 AT3 cells in 50μL Matrigel (Corning, NY, USA) were injected.

### Isolation of myeloid-derived suppressor cells (MDSCs)

MDSCs were isolated from spleens and tumors of tumor-bearing mice 3 weeks after orthotopic tumor cell implantation. Myeloid progenitor cells were isolated from the bone marrow of normal wild type mice. We used anti-Mouse Ly-6G, and Ly-6C antibody-conjugated magnetic beads (BD Biosciences, CA, USA). The purity of cell populations was > 99%. In short, the spleen was disrupted in PBS by using the plunger of a 3mL syringe, and then cell aggregates and debris were removed by passing cell suspension through a sterile 70μm mesh nylon strainer (Fisherbrand™). Mononuclear cells were separated by Lymphocyte separation medium (Corning®, NY, USA) as a white buffy coat layer. Cells were then centrifuged at 1500 rpm for 10 minutes followed by a washing step with PBS at 1200 rpm for 8 minutes. Then cells were resuspended at 1×10^8^ cells/mL in PBS and antibody conjugated with magnetic beads were added followed by incubation at 4°C for 30 minutes. Finally, positive cells were collected using a MACS LS column (Miltenyi Biotec, Germany) and a MidiMACS™ magnetic stand followed by a wash step with an extra amount of PBS. The purity of isolated MDSCs was checked by flow cytometry using Gr1 FITC and CD11b APC antibodies (purchased from Biolegend). MDSCs were grown in exosomes depleted media consisting of RPMI, 2mM L-glutamine, 1% MEM non-essential amino acids, 1mM sodium pyruvate and 10% FBS, supplemented with 100 ng/mL of GM-CSF.

### Exosome isolation

Exosomes were depleted from the complete media by ultracentrifugation for 70 minutes at 100,000x g using an ultracentrifuge (Beckman Coulter) and SW28 swinging-bucket rotor. The cells were then grown for 48 hours under normoxic condition. The cell culture supernatant was centrifuged at 700x g for 15 minutes to get rid of cell debris. To isolate exosomes, we employed combination of two steps of size-based method by passing through 0.20 µm syringe filter and centrifugation with 100k membrane tube at 3200x g for 30 minutes followed by a single step of ultracentrifugation at 100,000x g for 70 minutes (as described in our previous publication [15]).

### Protein quantification

Isolated exosomes resuspended in a minimal amount of PBS were lysed by RIPA buffer with protease and phosphatase inhibitor (100:1 dilution). Then exosomal protein was quantified by Bradford assay using Pierce™ BCA Protein Assay Kit (Thermo Scientific™) and serial dilution of BSA standard (Thermo Scientific™).

### Protein array

Proteins were extracted from tumor cells and their corresponding exosomes in both untreated and treated conditions to evaluate the expression profiles of 44 factors in duplicate by mouse cytokine antibody array (AAM-CYT-1000-8, RayBiotech, Inc.). 500μg of protein sample was loaded to the membrane according to the manufacturer’s instructions, and the chemiluminescent reaction was detected by using LAS-3000 imaging machine (Fuji Film, Japan). All signals (expression intensity) emitted from the membrane were normalized to the average of 6 positive control spots of the corresponding membrane using ImageJ software.

### *In vitro* migration assay

Trans-well assay was performed to evaluate the chemotaxis property of MDSC-derived exosomes. We used 24 trans-well plates with 8μm inserts in polyethylene terephthalate track-etched membranes (Corning, Inc.). We collected bone marrow cells and splenic mononuclear cells using Ficoll gradient centrifugation, and myeloid cells from bone marrow using CD11b+ magnetic beads from the normal Balb/c mice. 1.5×10^6^ cells/insert in serum-free media were added into the upper compartment of the chamber. Inserts will be placed in 12-well plates with DMEM containing 0.5% FBS in the presence or absence of exosomes isolated from MDSCs. After incubating overnight, migrated cells were collected from the bottom compartment for counting. insert membranes were washed, fixed and stained with 0.05% crystal violet to detect the migrated/invaded cells. The counting was made with an inverted microscope (Nikon Eclipse E200, Melville, NY, USA).

### *In vitro* scratch assay

Scratch assay was performed to detect the ability of MDSC-derived exosomes to increase migration and invasion of tumor cells. 4T1 luciferase positive cells were seeded in 6 well plates. After achieving 80±90% of confluency cells were starved overnight with 0.5% FBS for cell cycle synchronization and a measured wound was inflicted at the center of the culture (from top to bottom). Then, cells were treated with 100μl of splenic MDSC-derived exosomes in PBS containing 3×10^8^ exosomes for 48 hours in 2% FBS media. Microphotographs were taken every 24 hours using an automated all-in-one microscope (BZ-X710, Keyence). The wound size was measured using Image J software (NIH) by drawing a rectangular region of interest to quantify the visible area of the wound.

### Isolation of T-cells

Both CD4+ and CD8+ cells were isolated from normal mouse splenocytes by immune-magnetic negative selection kit (Stemcell Technology, Vancouver, Canada; Catalog #19852 and 19853, respectively). In short, harvested spleens from normal mice were disrupted in cold PBS containing 2% FBS. Clumps and debris were removed by passing the cell suspension through a 70-μm mesh nylon strainer. Then the single-cell suspension was centrifuged at 300 × g for 10 minutes and resuspended at 1×10^8^ nucleated cells/mL. Rat serum was added to the sample (50μL/mL) followed by the addition of isolation cocktail (50μL/mL). After mixing, the sample mix was incubated at room temperature for 10 minutes. RapidSpheres™ (75μL/mL) was added to the sample mix and incubated for 3 minutes. The tube was placed in an EASYSEP™ MAGNETS (Catalog #18001) for 3 minutes. The enriched cell suspension was collected by pouring off into a new tube. Cells were seeded in well plates bound with purified anti-mouse CD3e (5μL/mL) in T-cell media that consists of RPMI, 10% FBS, 1% MEAM, 2.5% HEPES, 1% penicillin-streptomycin, 0.5% β-mercaptoethanol, and purified anti-mouse CD28 (5μL/mL).

### Quantification of reactive oxygen species (ROS) generation by CD8+ T-cells

ROS production from CD8+ T-cells following MDSC-derived exosomes treatment in vitro was estimated by labeling the CD8+ T-cells using CM-H2DCFDA (Invitrogen™, C6827). In short CD8+ T-cells were isolated according to the above-mentioned method. After the final wash step of isolation, cells were resuspended in 1 mL PBS. DCFDA solution at a working concentration of 10 μM/mL was added followed by incubation in dark at 37°C for 30 minutes. Then the cells were washed with an extra amount of PBS to remove the unbound dye and resuspended with appropriate T-cell media. 100,000 cells were seeded per well of 96 well-plate. MDSC-derived exosomes were added in the treatment group and the same volume of PBS was added in the control group. Hydrogen peroxide (H2O2) was used as a positive control for ROS production. Following 4 hours of incubation, the fluorescent intensity of each condition was measured using Perkin Elmer Victor3 V 1420 multilabel plate reader, with excitation and emission wavelength of 485nm and 535nm, respectively. Each condition was triplicated for significance.

### CD8+ T-cell proliferation assay

Following isolation, 20,000 CD8+ cells were seeded in an anti-mouse CD3e-bound 96-well plate and treated with 10μL of MDSC-derived exosomes or the same volume of PBS (control). After 48 hours, 10μL of WST-1 reagent (Alkali Scientific Inc., FL, USA) was added to each well and incubated for 4 hours. The absorbance of each well was measured at a wavelength of 450 nm by the Perkin Elmer Victor3 V 1420 multilabel plate reader.

### Statistical analysis

Quantitative data were expressed as mean ± standard error of the mean (SEM) unless otherwise stated, and statistical differences between more than two groups were determined by analysis of variance (ANOVA) followed by multiple comparisons using Tukey’s multiple comparisons test. Comparison between 2 samples was performed by Student t test. GraphPad Prism version 8.2.1 for Windows (GraphPad Software, Inc., San Diego, CA) was used to perform the statistical analysis. Differences with p-values less than 0.05 were considered significant (*p<.05, **p<.01, ***p<.001, ****p<.0001).

## Results

### Isolation and characterization of exosomes from different MDSC population

We isolated MDSCs from normal (non-tumor bearing) wild-type mice bone marrow, and from spleen and tumor of tumor-bearing mice by magnetic particle separation using Ly6G and Ly6C beads. Tumors were implanted orthotopically in the mammary fat pad and allowed to grow for 3 weeks. Following the separation, by flowcytometry we estimated more than 98% of the cells are positive for MDSC markers (CD11b+Gr+) (**Figure 1A**). After 48 hours, we isolated exosomes from culture supernatant and characterized them by nanoparticle tracking analysis (NTA). Exosomes isolated from MDSCs of normal bone marrow (BM MDSC exo), the spleen of tumor-bearing mice (spleen MDSC exo) and tumor (tumor MDSC exo) were similar in size and distribution (**Figure 1B and 1C**). However, MDSCs from tumor released more exosomes compared to MDSCs from normal bone-marrow that could be due to the presence of the stressful condition, and more active and immunosuppressive MDSCs in primary tumor area (**Figure 1D**).

**Figure 1.**
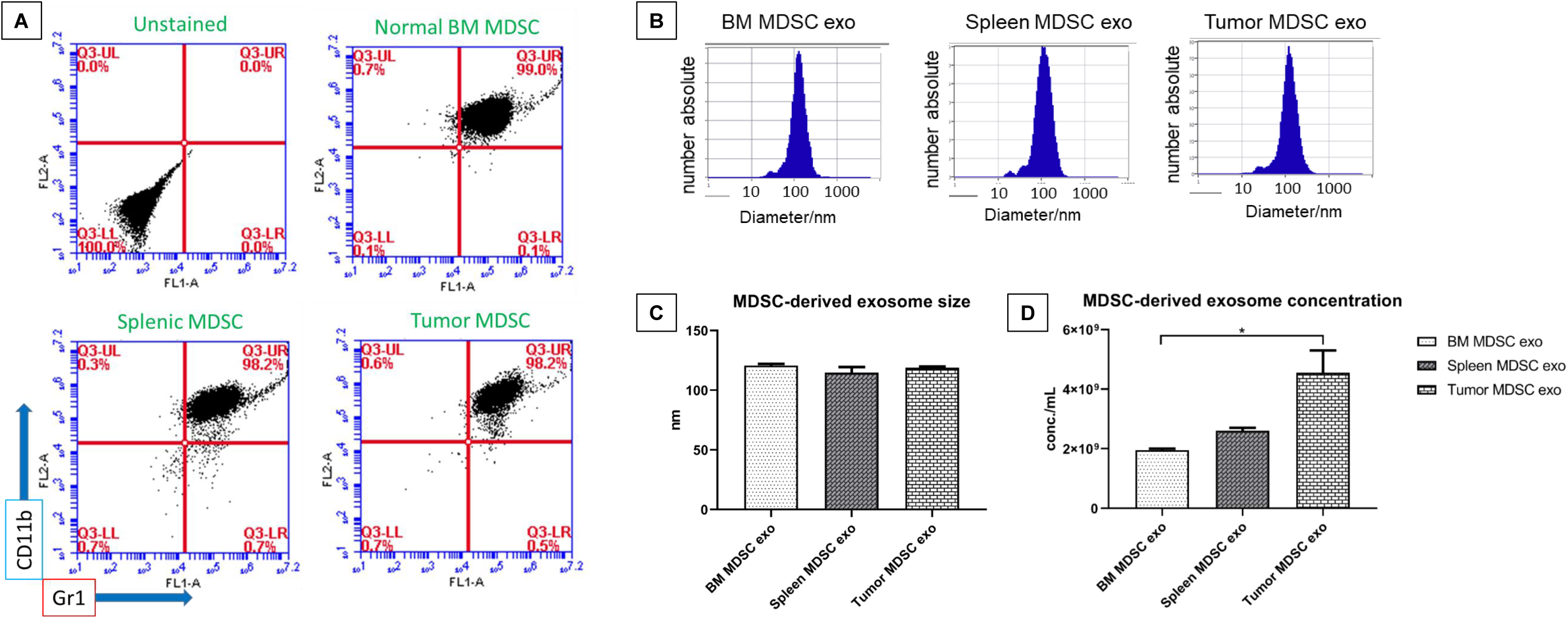
Isolation of MDSC-derived exosomes from different sources. **(A)** Flow cytometric analysis of isolated MDSCs from normal bone marrow (BM), the spleen of tumor-bearing mice and tumor, showing that more than 98% of cells are positive for CD11b and Gr1. **(B and C)** Nanoparticle tracking analysis showing no significant differences in the size distribution of exosomes isolated from MDSCs of normal bone marrow (BM), the spleen of tumor-bearing mice and tumor. **(D)** NTA analysis showing exosome concentration per mL. Quantitative data are expressed in mean ± SEM. *P<.05.

Next, we tested if protein contents of normal BM MDSC exo differ from the spleen MDSC exo and tumor MDSC exo isolated from tumor-bearing mice. We quantified the expression level of cytokines in MDSC-derived exosomes that are involved in tumor invasion (**Figure 2A**), angiogenesis (**Figure 2B**), and myeloid cell activation and function (**Figure 2C**) by membrane-based protein array. All the cytokines were significantly over-expressed in exosomes isolated from MDSCs of tumor-bearing mice (in both spleen MDSC exo and tumor MDSC exo) compared to normal BM MDSC exo. Interestingly, tumor MDSC exo showed a higher level of expression than that of spleen MDSC exo indicating that MDSC-derived exosomal cytokine contents are different based on the microenvironment of host tissues.

**Figure 2.**
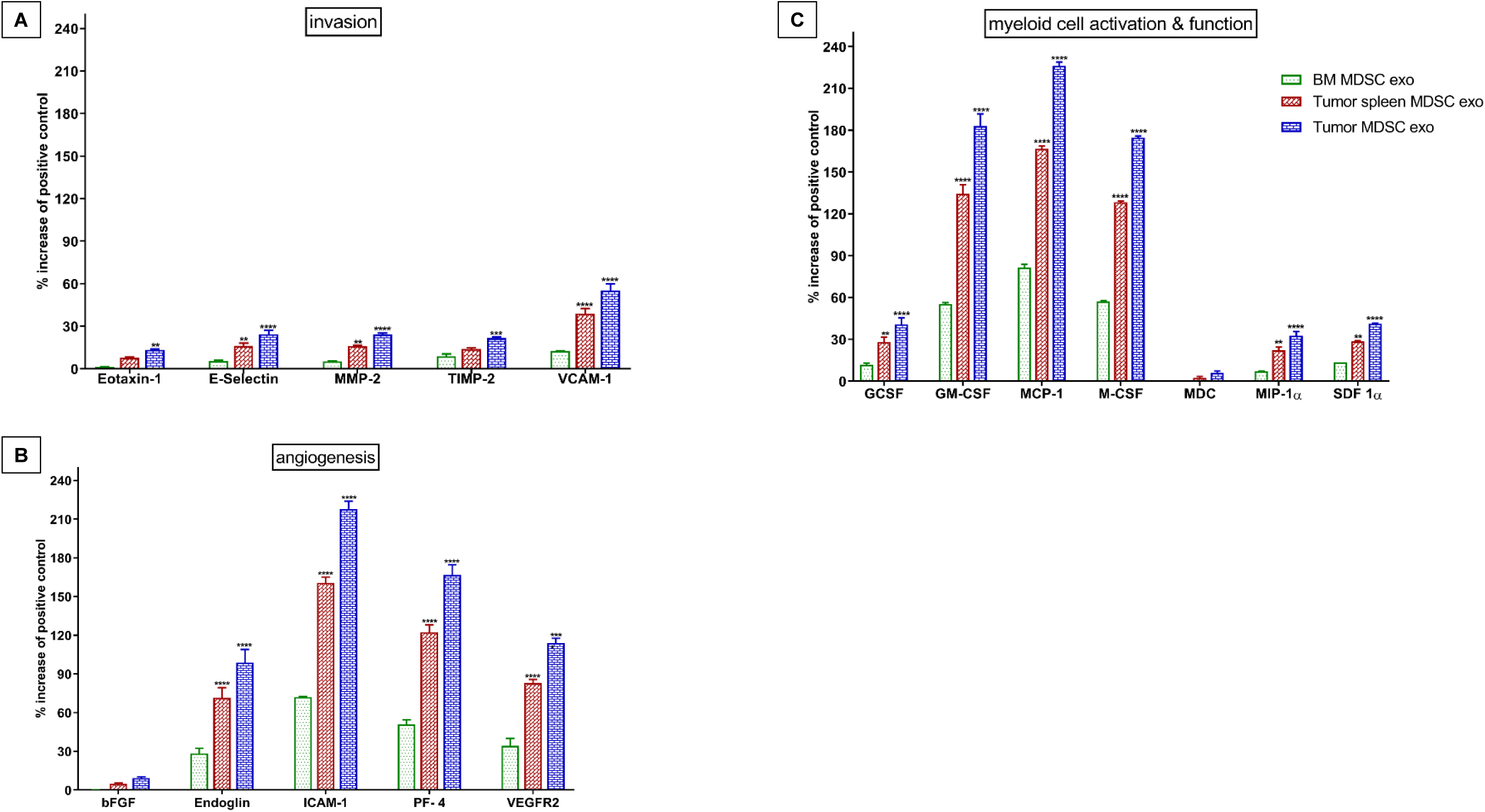
The expression level of cytokines in MDSC-derived exosomes that are involved in tumor invasion, angiogenesis, and myeloid cell activation and function. *In vitro* quantification of the level of cytokines associated with tumor **(A)** invasion, **(B)** angiogenesis, and **(C)** myeloid cell activation and function, detected in the membrane-based array in protein samples collected from the exosomes isolated from MDSCs of normal bone marrow (BM), spleen of tumor-bearing mice and tumor. Quantitative data are expressed in mean ± SEM. *P<.05, **P<.01, ***P<.001, ****P<.0001. n = 4.

### MDSC-derived exosomes promote invasion and migration of tumor cells

MDSCs were demonstrated to promote tumor invasion and metastasis by two mechanisms: (i) increased production of multiple matrix metalloproteinases (MMPs) for extra-cellular matrix degradation, and chemokines to establish a pre-metastatic milieu [16, 17], and (ii) merging with tumor cells, MDSCs promoting the metastatic process [18, 19]. We observed significantly higher expression of invasion and migration-associated cytokines in the spleen MDSC-exo of tumor-bearing mice compared to normal BM MDSC-exo, which led us to further investigate the role of MDSC-derived exosomes in promoting invasion and migration of tumor cells. *In vitro* wound-healing assay showed a significant increase of invasion and migration of 4T1 tumor cells in spleen MDSC-exo treated group compared to the untreated control group in 24 and 48 hours (**Figure 3A and 3B**).

**Figure 3.**
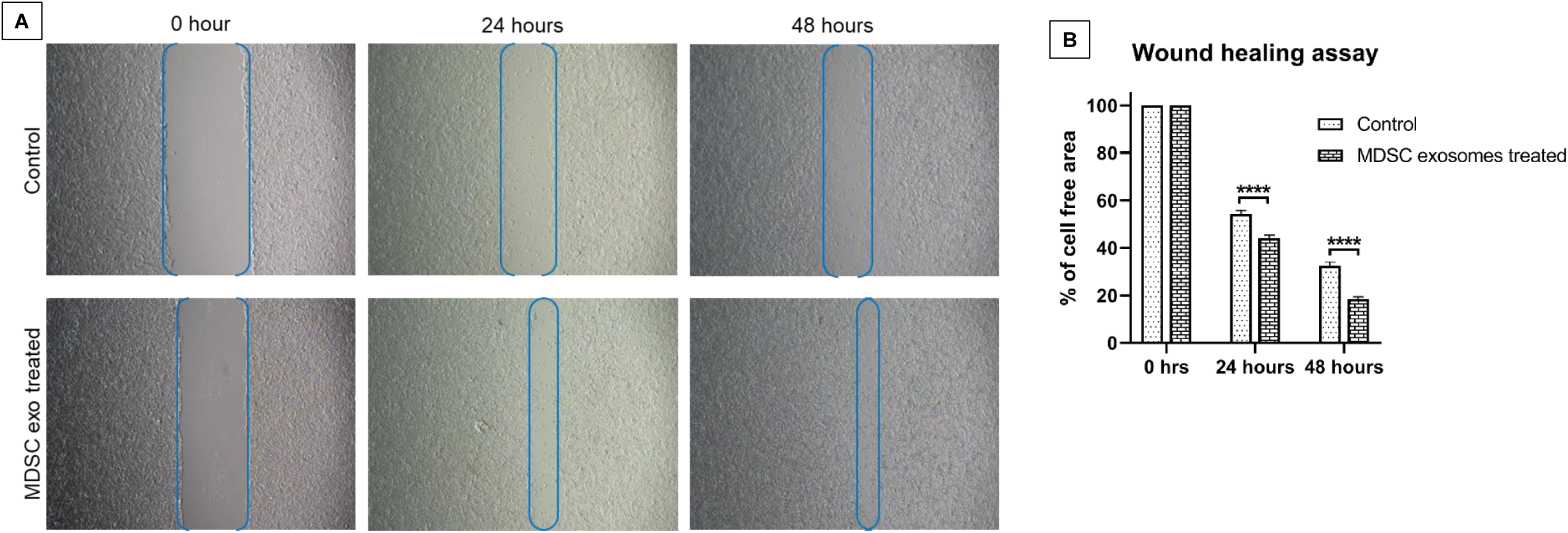
Role of MDSC-derived exosomes in tumor cell migration. A wound-healing assay/scratch assay was carried out in 4T1 murine breast cancer cell line with MDSC-derived exosomes treatment or without treatment (control). **(A)** Representative microscopic images (4x) are shown before treatment, and 24 and 48 hours after treatment. **(B)** Semi-quantitative analysis of the percentage of non-covered area/cell-free area. Quantitative data are expressed in mean ± SEM., ****P<.0001. n = 10.

### MDSC-derived exosomes promote the recruitment of immunosuppressive cells *in vitro*

Tumor-specific endocrine factors systemically stimulate the quiescent immune-compartments (bone marrow, spleen, lymph nodes), resulting in the expansion, mobilization, and recruitment of immunosuppressive cells. Discrete subsets of tumor-instigated immune cells bolster tumor progression and metastasis by governing angiogenesis, inflammation and immune suppression. Of the immune cells, much focus has been denoted towards the MDSCs [20], tumor associated macrophages (TAMs) [21], Tie2-expressing monocytes [22], vascular endothelial (VE)-cadherin+CD45+ vascular leukocytes, and infiltrating mast cells and neutrophils [23, 24]. We observed significant high expression levels of cytokines crucial for immunosuppressive cell mobilization and recruitment in spleen MDSC-exo isolated from splenic MDSCs of tumor-bearing mice compared to normal BM MDSC exo.

To determine the chemotaxis capability of MDSC exo, we seeded CD11b+ myeloid cells (from bone marrow) or all bone marrow cells, or all splenic mononuclear cells (following ficoll separation) isolated from normal mice in the upper chamber and with or without spleen MDSC exo in the bottom chamber of the trans-well insert. After 24 hours of incubation, the number of migrated cells in the bottom chamber was significantly higher in the wells treated with MDSC exo compared to untreated control wells (**Figure 4A, 4B and 4C**). We also washed, fixed and stained the insert membranes (of CD11b+ cells incubated group) with 0.05% crystal violet to detect the migrated/invaded cells. The number of cells that are attached to the membrane was visualized by microscopy (**Figure 4D**) and later quantified by ImageJ cell counter. A significantly higher number of CD11b+ myeloid cells were attached to the transmembrane in the wells treated with spleen MDSC exo compared to untreated control wells (**Figure 4E**).

**Figure 4.**
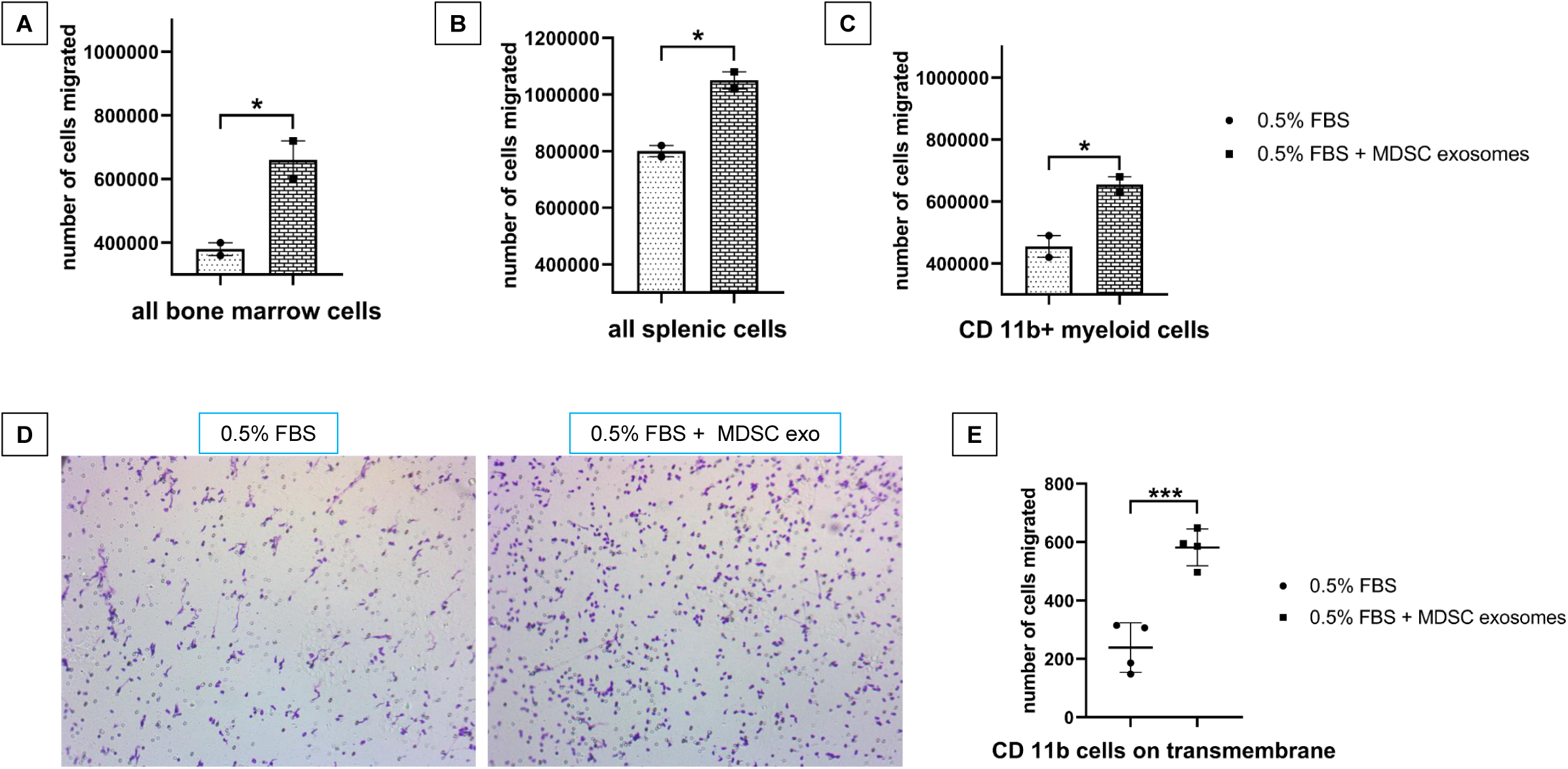
Role of MDSC-derived exosomes in immune cell migration. Isolated mouse myeloid cells, bone marrow cells and splenic cells were seeded on the top chamber of transwell, and MDSC-derived exosomes were added in the bottom chamber with 0.5% FBS. After 24 hours, migrated **(A)** bone marrow cells, **(B)** splenic cells and **(C)** myeloid cells in the bottom chamber were counted. In addition to that after 24 hours, **(D)** attached myeloid cells on the transwell membrane were visualized under the light microscope, and **(E)** were quantified. Quantitative data are expressed in mean ± SEM. *P<.05, ***P<.001, n = 4.

### Expression of T-cell function-associated and immunomodulatory cytokines in exosomes from different MDSC population

We further estimated the level of expression of T-cell function-associated and immunomodulatory cytokines in protein contents of normal BM MDSC exo and exosomes isolated from MDSCs (tumor and splenic) of tumor-bearing mice by protein array. Among immunomodulatory cytokines level of IL-12, IL-13, IL-1ra, IL-4, C-X-C motif chemokine 5 (LIX) and TNF-α were significantly elevated in tumor-MDSC-exo and spleen-MDSC-exo of tumor-bearing mice compared to normal BM-MDSC-exo (**Figure 5A**). Among T-cell function-associated cytokines, IL-2, IL-7, L-selectin and thymic stromal lymphopoietin (TSLP) were significantly high in exosomes derived from tumor or splenic MDSC of tumor-bearing mice compared to normal BM MDSC-exo (**Figure 5B**).

**Figure 5.**
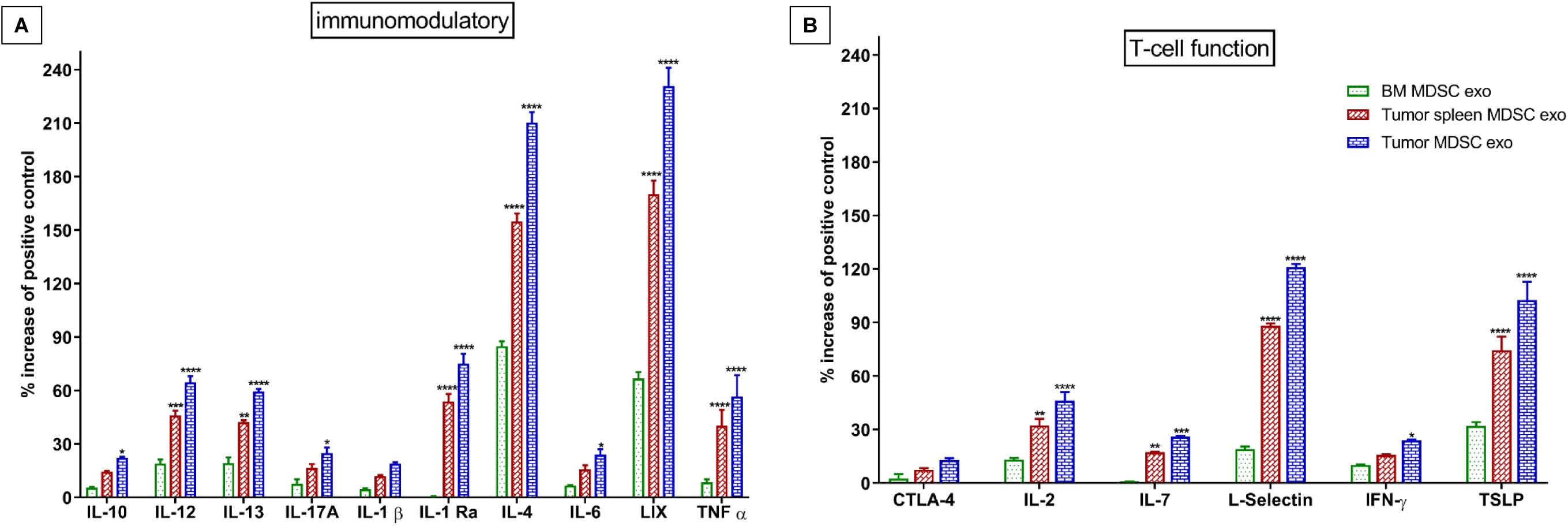
Expression level of cytokines in MDSC-derived exosomes that are involved in T-cell function and immunomodulation. *In vitro* quantification of the level of cytokines associated with (A) immunomodulation and (B) T-cell function, detected in the membrane-based array in protein samples collected from the exosomes isolated from MDSCs of normal bone marrow (BM), the spleen of tumor-bearing mice and tumor. Quantitative data are expressed in mean ± SEM. *P<.05, **P<.01, ***P<.001, ****P<.0001. n = 4.

### *In vivo* effect of MDSC-derived exosomes on T-cells

Next, we investigated whether MDSC-derived exosomes treatment could deplete the CD8+ T-cells in mice. For this *in vivo* study, we used both C57BL/6 and Balb/c normal mice. We treated the mice with MDSC-derived exosomes by IV injection through the tail vein for a week (total of 3 doses, alternative day). Then we euthanized the animals and harvested spleens for flow-cytometric evaluation. CD8+ T-cell population in splenic MDSC-exosomes treated animals was remarkably declined compared to the untreated control group in both animal models (**Figure 6A and 6B**). However, we did not observe any significant change in CD4+ T-cell population.

**Figure 6.**
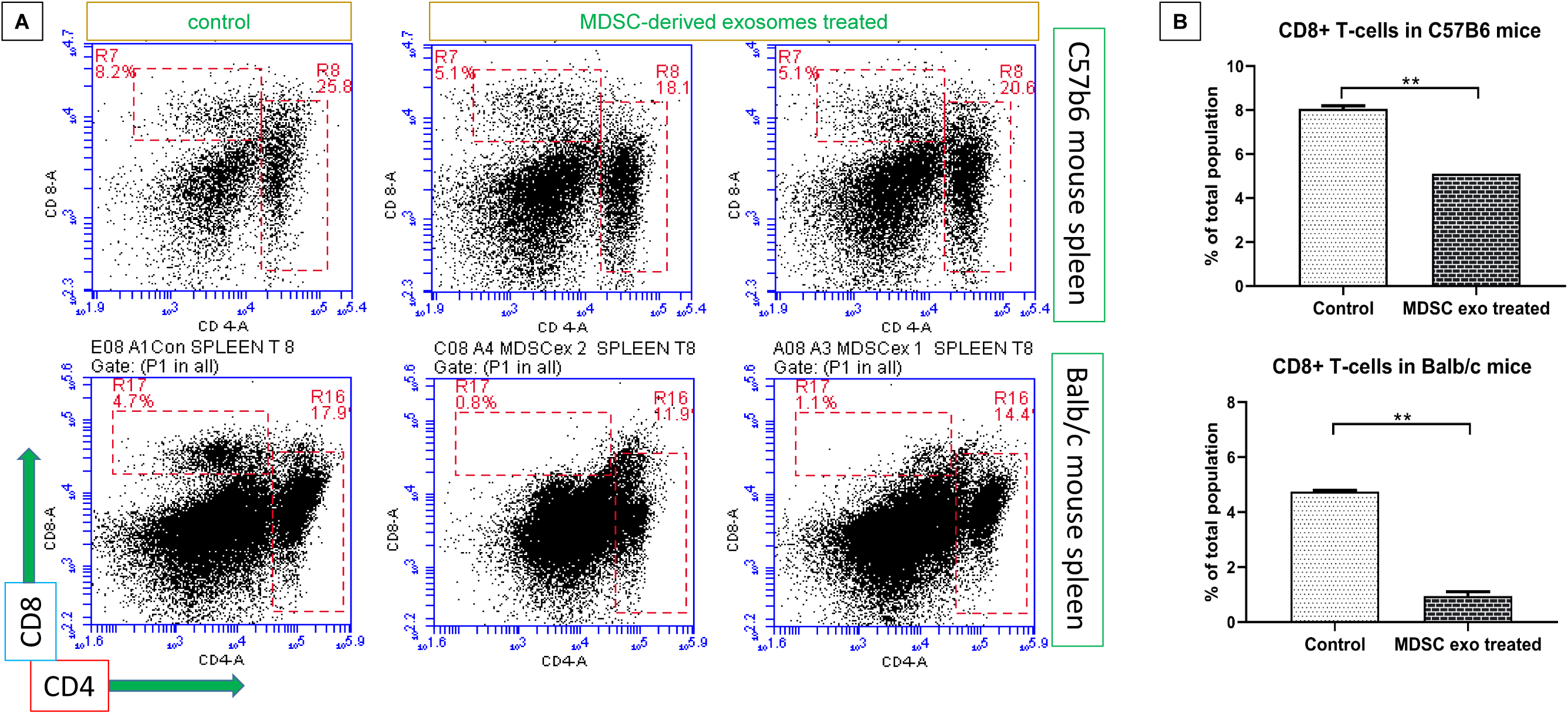
*In vivo* depletion of CD8+ T-cells by MDSC-derived exosomes treatment. Normal Balb/c and C57BL/6 mice were treated with MDSC-derived exosomes for 1 week and without treatment (control). (A) Representative flow-cytometric plots and (B) quantification of cells from the spleen showing decreased CD8+ T-cells. Quantitative data are expressed in mean ± SEM. **P<.01, n = 4.

### In vivo effect of MDSC-derived exosomes on myeloid cells

We also explored if the MDSC-derived exosomes treatment could change the distribution of the myeloid populations *in vivo*. We noticed a significant reduction of M1-macrophages (CD11b+CD80+ and CD11b+CD86+) in the spleen of whereas no notable changes in the other organs (**Figure 7A)**. There was a considerable decline of the monocytic MDSCs (CD11b+Gr1+Ly6C+) and the expansion of granulocytic MDSCs (CD11b+Gr1+Ly6G+) in the spleen of the treated animals compared to the untreated group (**Figure 7B)**.

**Figure 7.**
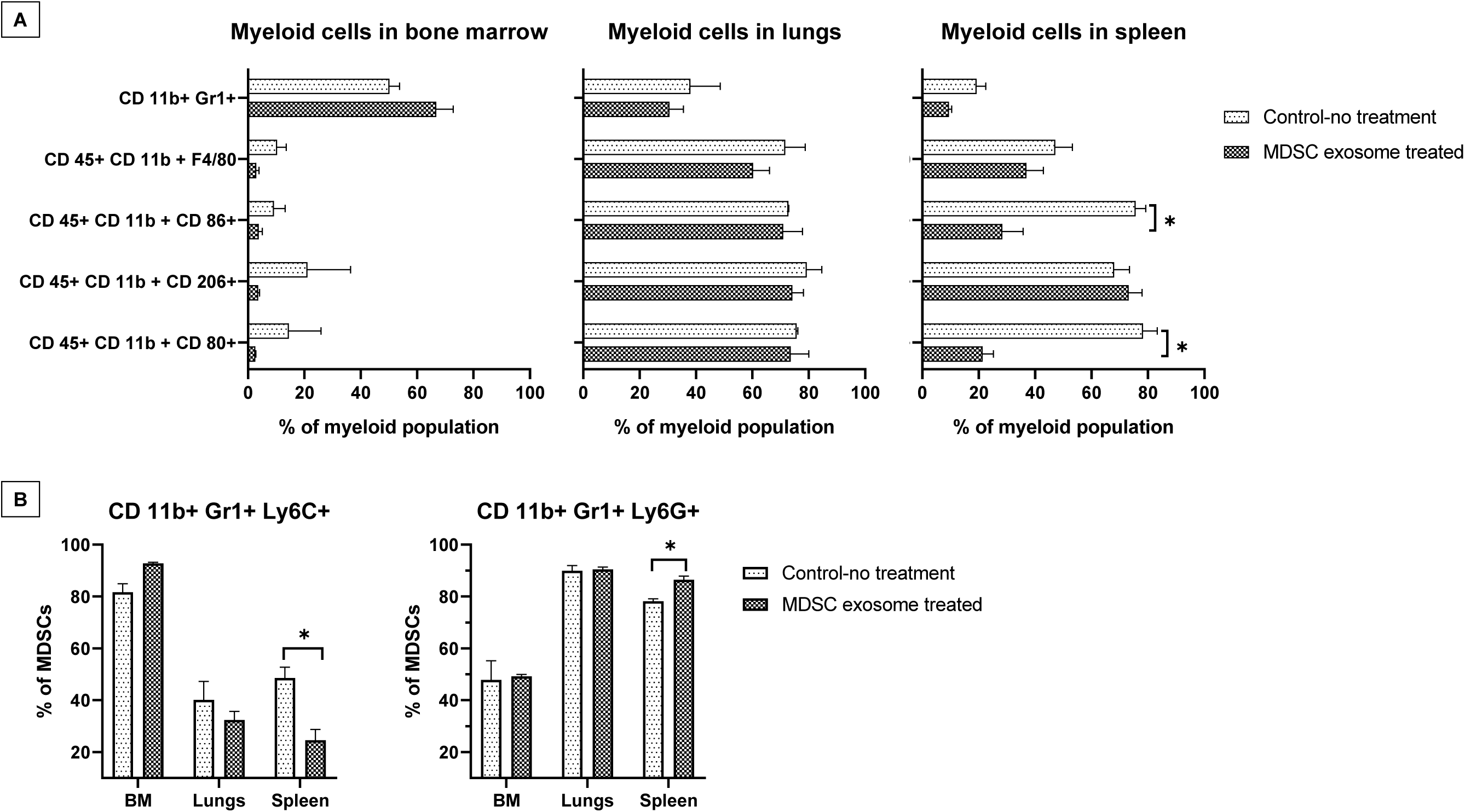
*In vivo* depletion of M1-macrophages and increased gMDSCs by MDSC-derived exosomes treatment. Normal Balb/c mice were treated with MDSC-derived exosomes for 1 week and without treatment (control). Quantification of cells from the bone marrow, lungs, and spleen showing **(A)** decreased M1-macrophages in spleen and **(B)** decreased monocytic MDSCs and increased granulocytic MDSCs in spleen. Quantitative data are expressed in mean ± SEM. *P<.05, n = 3.

### *In vitro* effect of MDSC-derived exosomes on T-cells

As we have seen, MDSC exo express a significantly high level of cytokines that facilitate regulatory T-cell or Th2 cell functions and immunosuppression, we wanted to investigate the effect of MDSC exo directly on CD4+ and CD8+ T-cells *in vitro*. We isolated both cells and treated them with MDSC exo or with the same volume of PBS (control). After 24 hours, we collected the cells and analyzed the functional marker changes by flow-cytometry. Splenic MDSC exo treated CD4+ T-cells expressed a significantly higher level of T-regulatory cell marker (CD25) and Th2 cell marker (CD184) compared to the control group (**Figure 8A**). There was no change in the level of T-cell activation or exhaustion marker (CD279/PD-1). For the CD8+ T-cells, MDSC exo treated cells showed a significantly higher level of T-cell activation marker (CD44), naïve T-cell marker (CD62L), and exhaustion marker (CD279) (**Figure 8B**).

**Figure 8.**
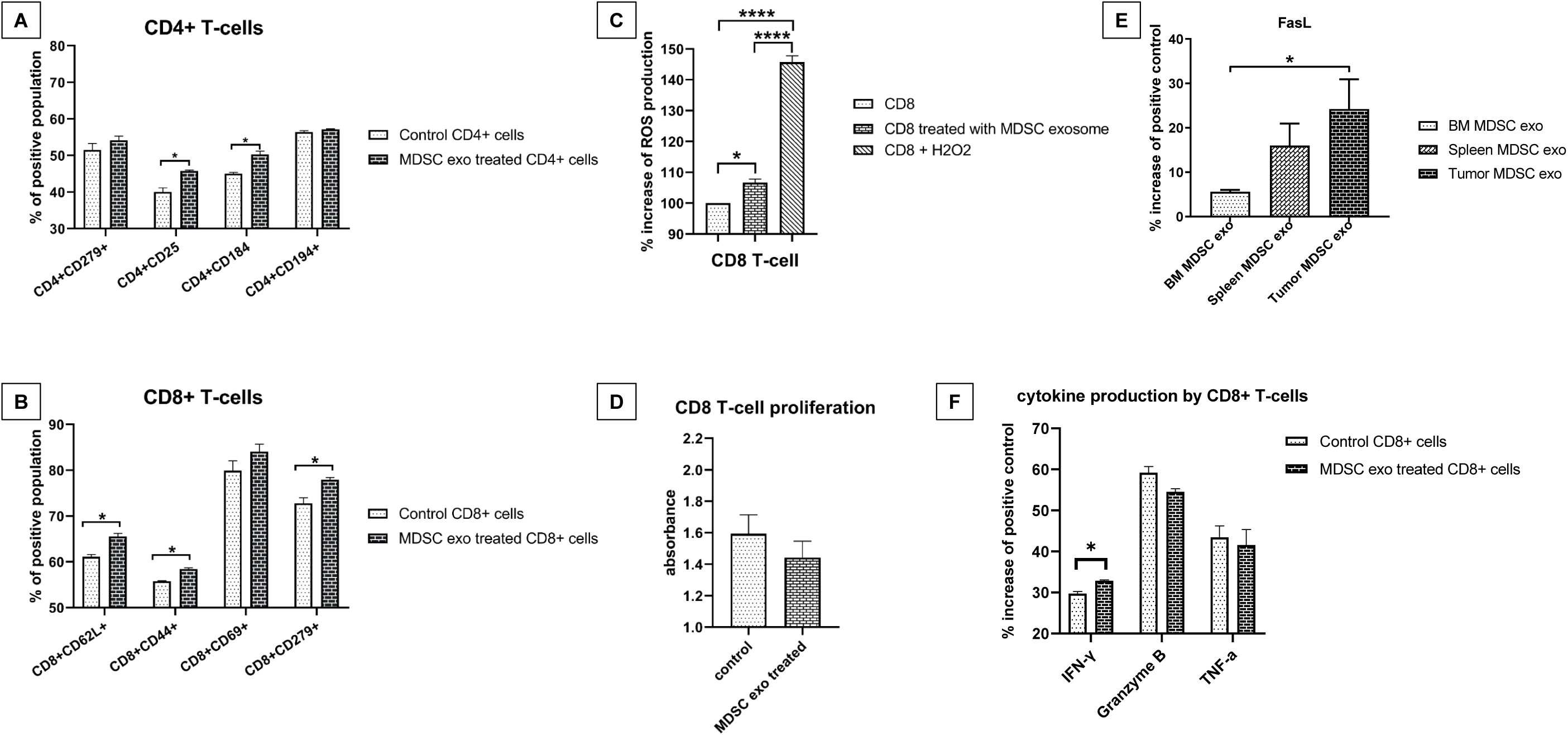
Effect of MDSC-derived exosomes on CD4 and CD8-positive T-cells *in vitro*. **(A and B)** Isolated CD4 and CD8-positive T-cells were co-cultured with or without MDSC-derived exosomes treatment for 24 hours followed by flow-cytometric analysis of the cells. **(C)** Effect of MDSC-derived exosomes on ROS production by CD8+ T-cells was determined by CM-H2DCFDA-labeled CD8+ positive T-cells treated with MDSC-derived. **(D)** Effect of MDSC-derived exosomes on CD8+ positive T-cell proliferation by a cell proliferation assay using WST-1 reagent. Membrane-based protein array was used to determine the **(E)** level of the FasL in exosomes from different MDSC populations and **(F)** expression levels of cytokines in CD8 T-cells. Quantitative data are expressed in mean ± SEM. *P<.05, ****P<.0001. n = 4.

### *In vitro* effect of MDSC-derived exosomes on CD8+ T-cells function and proliferation

MDSCs release ROS molecules as part of a primary mechanism to suppress T cell responses [25]. Considering the previous results on CD8+ T-cell, we further determined the level of ROS production by CD8+ T-cell after splenic MDSC exo treatment and also if the treatment affects CD8+ T-cell proliferation. We labeled the CD8+ T-cells with CM-H2DCFDA ROS probe and treated them with MDSC exo or with PBS (control). After 4 hours, MDSC exo treated CD8+ T-cells showed a considerably higher amount of ROS production compared to the control (**Figure 8C**). Although the cell proliferation assay using WST-1 reagent showed a decreased number of CD8+ T-cells in the MDSC exo group compared to the control group, it was not statistically significant (**Figure 8D**).

Considering the fact that MDSC exo were able to activate and deplete CD8+ T-cells, we determined the level of FasL in different MDSC exo that could conceivably trigger the apoptosis process in CD8+ T-cells. We detected higher expression of FasL in the exosomes isolated from the MDSCs in tumors compared to that in exosomes of MDSCs from spleen and bone marrow (**Figure 8E**). Furthermore, we quantified the activation markers of CD8+ T-cells with or without treatment with MDSC exo by protein array. Treatment with MDSC exo appreciably increased the level of IFN-γ while no significant changes in Granzyme B and TNF-α level (**Figure 8F**).

## Discussion

It has been perceived that functional differences may exist in MDSCs isolated from different environments within the same host, and MDSCs from tumors have a stronger immunosuppressive capacity than MDSCs in the peripheral lymphoid organs (spleen, lymph nodes) [26]. We observed that exosomes were secreted more abundantly from tumor MDSCs, presumably due to the fact that MDSCs in the TME are in a more distressing milieu (hypoxia, acidic pH, etc.). It has been contemplated that cells may exploit exosome secretion to survive under stressful conditions [27-29]. We also observed a higher concentration of proteins that are crucial for tumor growth, invasion, angiogenesis, and immunomodulation in exosomes isolated from MDSCs in tumor than those are in the spleen or bone marrow. Although Haverkamp et al. reported that MDSCs from inflammatory sites or from tumor tissue possess the immediate ability to hamper T-cell function, whereas those isolated from peripheral tissues were not suppressive without activation of iNOS by exposure to IFN-γ [30], we noticed equivalent competency in MDSC exo isolated both from tumor and spleen. However splenic MDSC-derived exosomes demonstrated that to a lesser extent.

MDSCs are competent in promoting tumor growth through remodeling the TME [19, 31]. But then again MDSCs are a miscellaneous population of the immature myeloid cells that comprise of monocytic and granulocytic subpopulations both of which have been shown to be immunosuppressive [32]. It has been recently reported that early expansion and infiltrations of mMDSCs take place in the primary tumors where they pave the way for tumor cell dissemination by inducing epithelial to mesenchymal transition (EMT), and higher levels of gMDSCs infiltrations in the metastatic site where they augment the colonization and metastatic growth of disseminated tumor cell by reverting EMT [33]. We observed that treatment of normal wild type mice with MDSC exo significantly decrease mMDSCs and increase gMDSCs in the spleen. As expected, we also detected a decrease in M1-macrophages in the spleen.

We observed that MDSC-derived exosomes are able to deplete CD8+ T-cells *in vivo* and inhibit the proliferation of CD8+ T-cells *in vitro*. When activated through their antigen-specific receptor (TCR) and CD28 co-receptor, resting mature T lymphocytes start to proliferate followed by the so-called activation-induced cell death (AICD), which mechanistically is triggered by the death receptor and leads to apoptosis. The apoptotic pathway is triggered by signals originating from cell-surface death receptors that are activated by several ligands such as CD95L (FasL), TNF or TNF-related apoptosis-inducing ligand (TRAIL) [34]. Our protein array data demonstrated that MDSC-derived exosomes contain a high level of Fas and TNF-1α. We noticed that MDSC-derived exosome treatment increases the activation markers of CD8+ T cells (CD69 and CD44), as well as their exhaustion marker CD279/PD-1. CD8+ T-cells also kill target cells by a cytokine-mediated mechanism (IFN-γ, TNF-α, etc), which are produced and secreted as long as TCR stimulation continues. IFN-γ induces transcriptional activation of the MHC class I antigen presentation pathway and Fas in target cells, leading to enhanced Fas-mediated target-cell lysis [35]. We noted that MDSC exo can activate CD8+ T cells and prompt them to generate more IFN-γ. Interestingly, ROS can control the fate of antigen-specific T cells through reciprocal modulation of the main effector molecules FasL and Bcl-2 [36]. We detected a significantly large amount of ROS production from the CD8+ T-cells that were treated with MDSC-derived exosomes. Therefore, we hypothesized that MDSC-derived exosomes precipitate CD8+ T-cell apoptosis by AICD through hyper-activation or repeated stimulation, which in turn results in increased levels of ROS production and activation of Fas/FasL (CD95/CD95L) pathway.

In summary **(Figure 9)**, we comprehensively demonstrated that MDSC-derived exosomes inherit pro-tumorigenic factors and functionally resemble parental cells in immunosuppression, tumor growth, angiogenesis, invasion and metastasis. In addition, MDSC-derived exosomes are capable of increasing ROS production and inciting Fas/FasL pathway in CD8+ T-cells that precipitate AICD.

**Figure 9.**
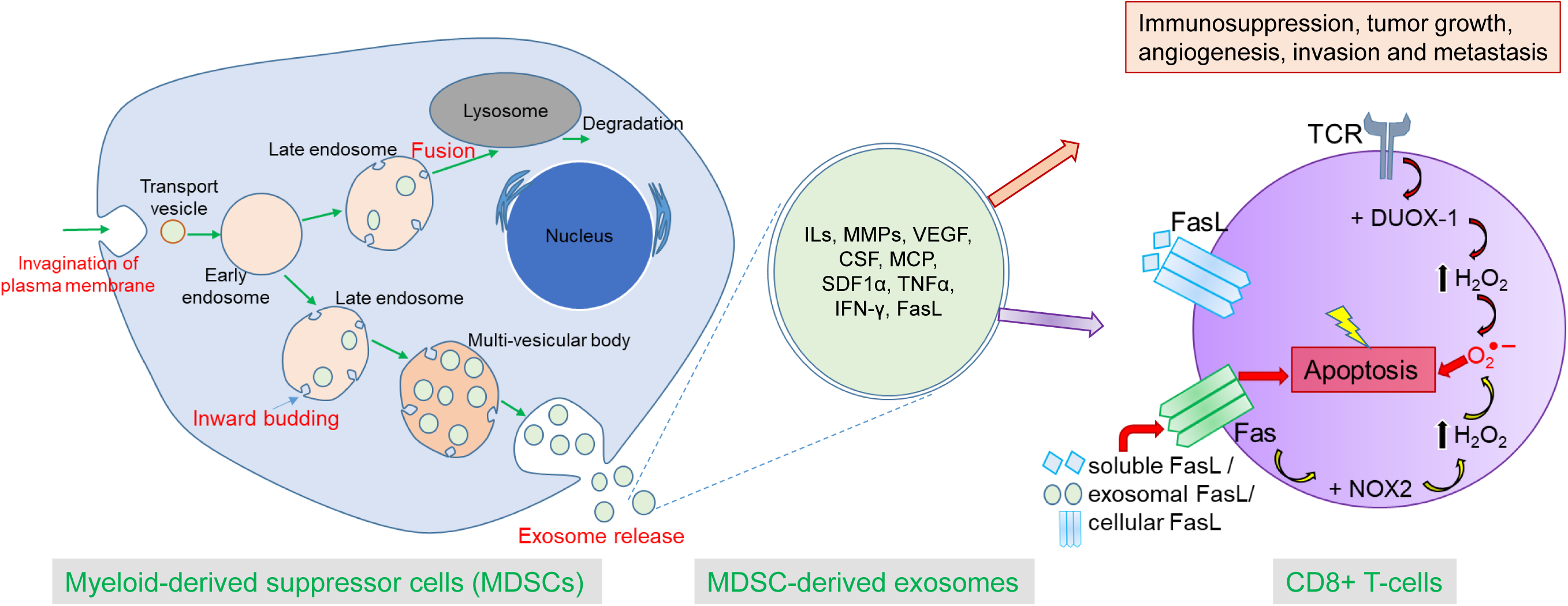
Schematic diagram showing process of biogenesis of exosomes from MDSCs and role of MDSC-derived exosomes in tumor progression and immunosuppression by AICD. Exosmes secreted from the MDSCs contain pro-tumorigenic factors from the parent cells and can play a crucial role in immunosuppression, tumor growth, angiogenesis, invasion and metastasis by dispensing their contents to the other TME cells or distant cells. MDSC-derived exosomes can activate CD8+ T-cells and TCR triggering causes activation of DUOX-1 that leads to H_2_O_2_ production and eventually generation of ROS in mitochondria. Prolonged TCR stimulation triggers overexpression of both Fas (receptor) and FasL (ligand) which culminates in fratricide (from direct cell contact) or autocrine suicide (interaction of soluble FasL with Fas).

## Author Contributions

<u>Mohammad Harun Rashid:</u> Conceived the hypothesis, designing and performing the experiments, data collection, data analysis and interpretation, and wrote the manuscript.

<u>Thaiz F Borin:</u> acquisition of *in vitro* data and edited the manuscript.

<u>Roxan Ara:</u> helped with treating the animals.

<u>Raziye Piranlioglu:</u> edited the manuscript.

<u>Bhagelu R Achyut:</u> helped with planning *in vitro* T-cell experiments.

<u>Hasan Korkaya:</u> provided the guidance for MDSC collection and types, and help implantation breast cancers.

<u>Yutao Liu:</u> provided lab facilities for NTA, data interpretation and editing of the manuscript.

<u>Ali S Arbab</u>: supervised the findings of this work, aided in interpreting the results, provided the funds and critical revision of the manuscript.

